## Supplementary Information for "Systematic exploration of bacterial form I rubisco maximal carboxylation rates"

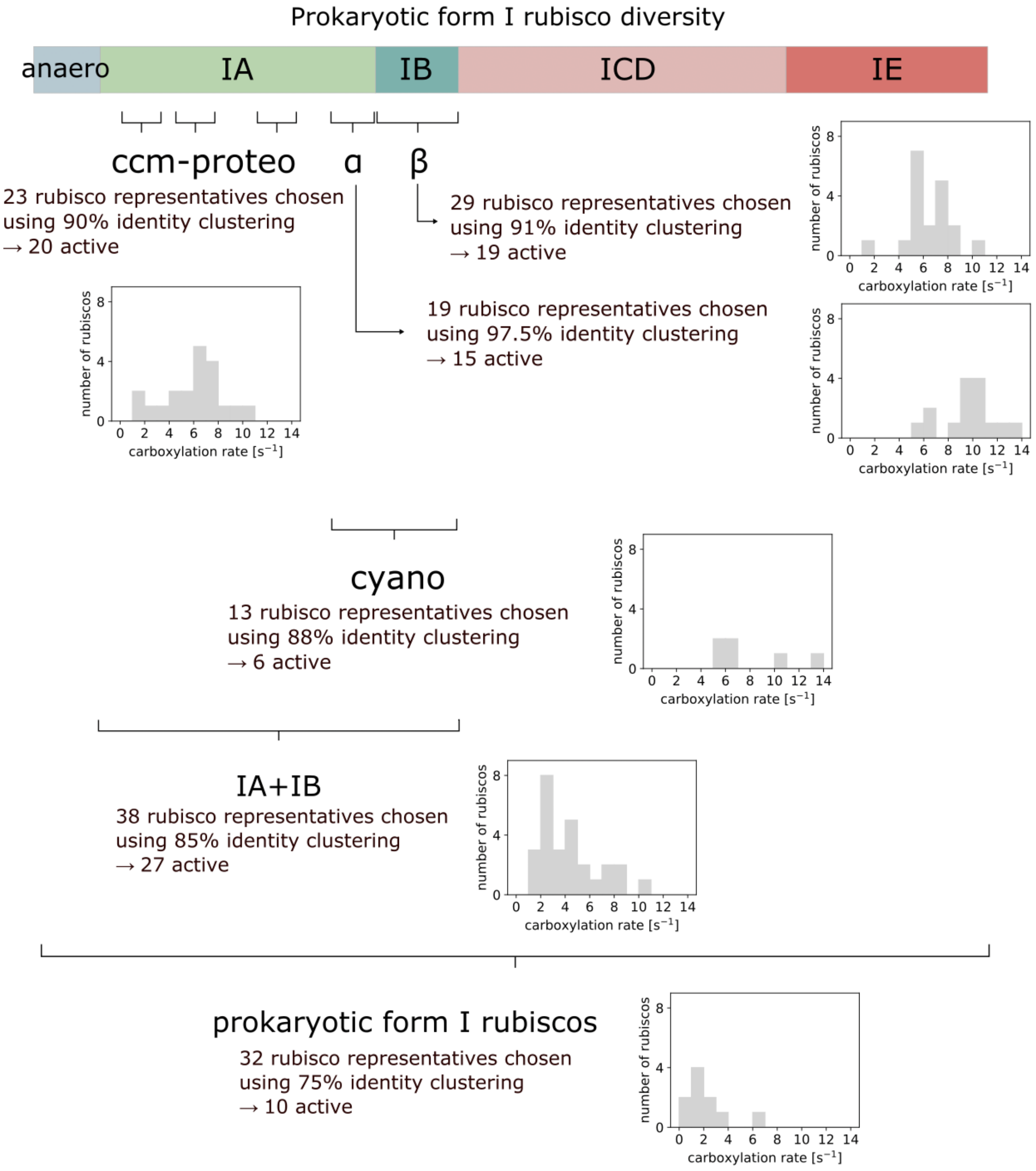


***Supplementary Fig. 1. Sequential screening strategy used for the selection of rubisco variants for characterization.*** *An iterative clustering approach was employed to screen the totality and specific subgroups of the form I rubisco family. The number of active rubiscos and measured carboxylation rates are indicated at each step.*

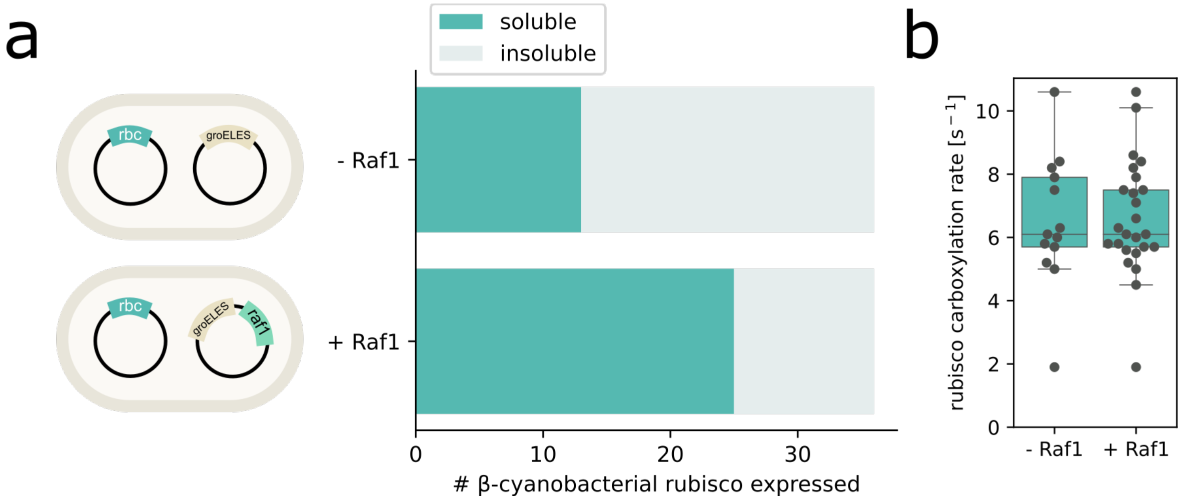


***Supplementary Fig. 2. Co-expression of β-cyanobacteria rubisco with Raf1 from Euhalothece natronophila greatly enhances solubility.*** *(A) rbcL-rbcX-rbcS operon containing plasmid was cotransformed with a plasmid expressing groEL-groES with or without raf1 in E. coli. The presence of Raf1 almost doubled the number of soluble and active rubiscos among the 36 tested β-cyanobacteria variants. (B) The average rate of β-cyanobacterial rubiscos is not changed by adding the newly soluble rubiscos.*

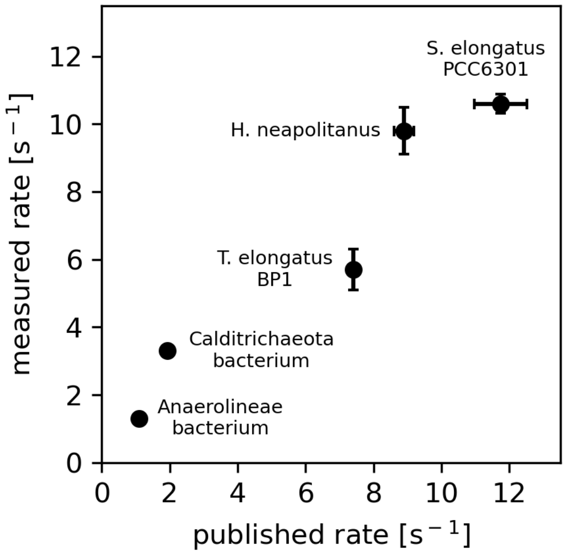


***Supplementary Fig. 3. Carboxylation rates measured in this study have similar ranking as rates published in the literature in spite of different techniques and conditions.*** *Values in both axes are not supposed to be equal as those measured in this study result from coupled assays, which tend to underestimate the rates compared to direct assays used in the literature, and on the other side they were performed at 30°C, resulting in faster rates compared to literature measurements done at 25°C.*

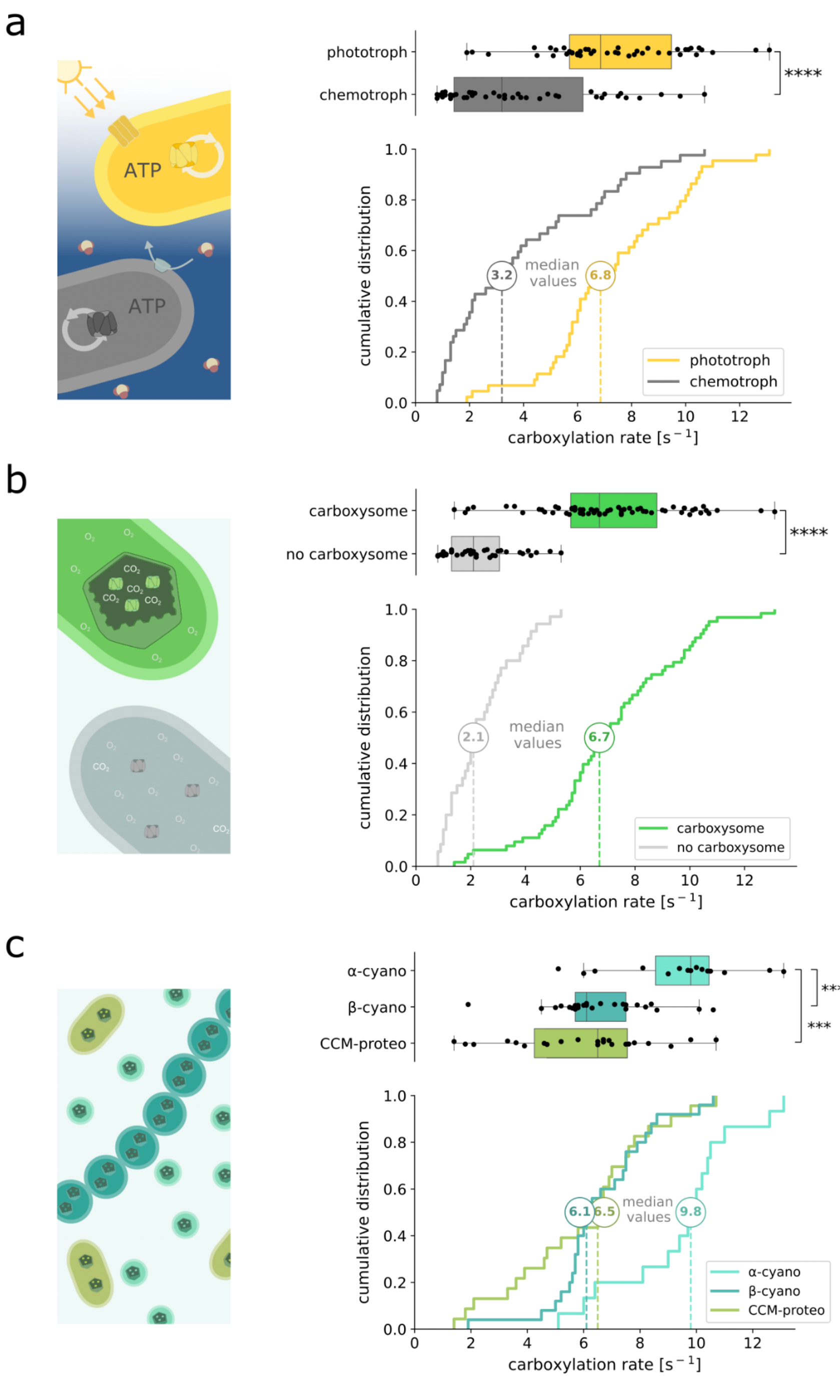


***Supplementary Fig. 4. Comparative analysis of all measured rubisco rates from this study in function of their associated biological parameters.*** *Box and cumulative distribution plots of all rubisco carboxylation rates from this study from different clusters: chemo- and phototrophic bacterial rubiscos (A), carboxysome-associated rubiscos and their counterparts (B), α- and β- cyanobacterial, and carboxysome-associated proteobacterial rubiscos (C). Contrary to Fig. 2 where uniform sampling was ensured, present plots contain all rates measured in this study. Mann-Whitney U test (A and B) or Kruskal-Wallis followed by Dunn multiple comparison tests (C) were applied. ***p < 0.001, ****p < 0.0001. Legend abbreviations are as follows: α-cyano, α-cyanobacterial rubisco; β-cyano, β-cyanobacterial rubisco; CCM-proteo, carboxysome-associated proteobacterial rubisco.*

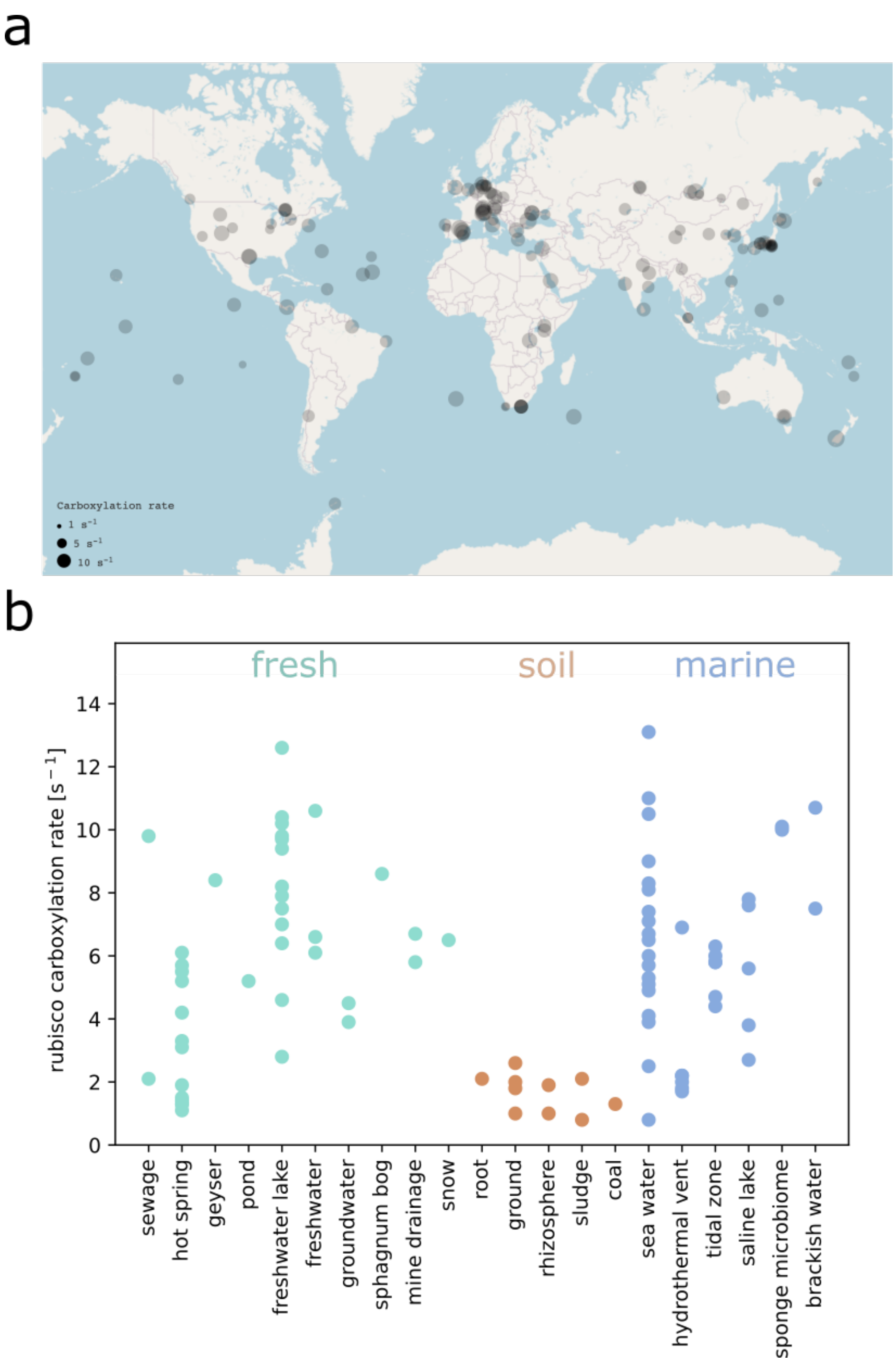


***Supplementary Fig. 5. Environmental context of rubisco sequences tested in this study.*** *A) Map of rubisco carboxylation rates measured in this study, at the GPS coordinates of the biological samples associated with the cognate rubisco genes. Dots' area is proportional to measured carboxylation rates in vitro. B) Rubisco carboxylation rate in function of its bacteria habitat. Environmental information of the biological sample containing each rubisco's sequence was collected and sorted into main habitats.*

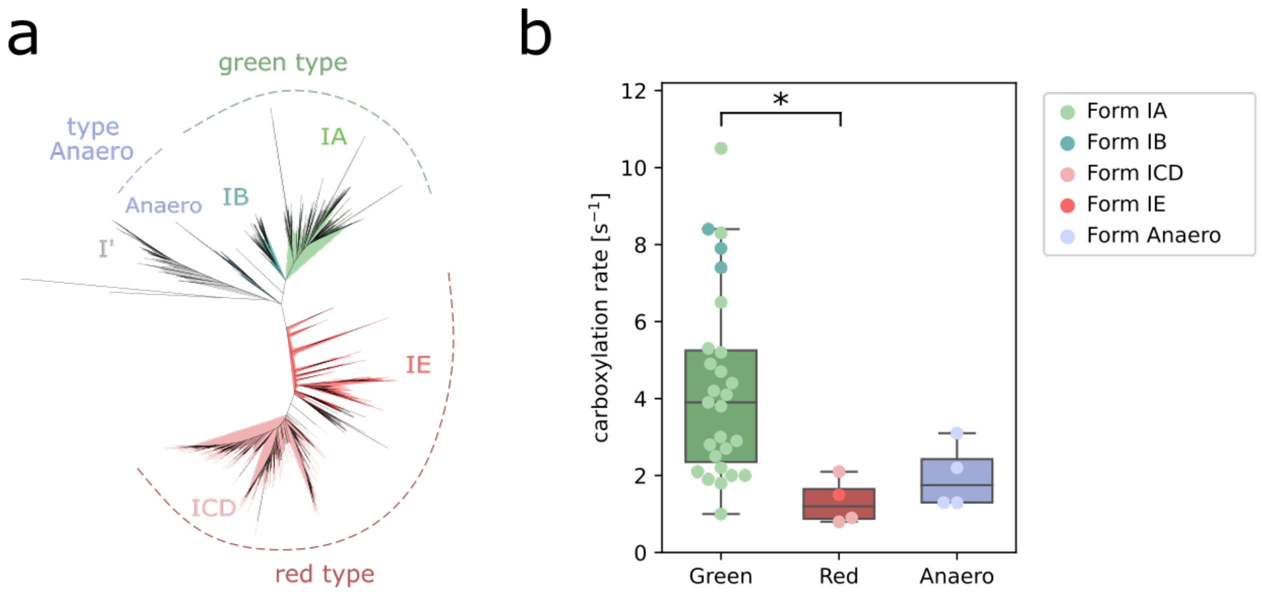


***Supplementary Fig. 6. Green type rubiscos are faster than red type ones.*** *A) Form I rubisco large subunit phylogenetic tree showing the 3 distinct types: the "green", the "red", and the newly discovered "anaero" types (1). B) Box plot of carboxylation rates from rubiscos of the 3 types. To ensure unbiased study of each group, we selected a set of rubiscos uniformly covering their genetic diversity. Kruskal-Wallis followed by Dunn multiple comparison tests were applied. *p < 0.05.*

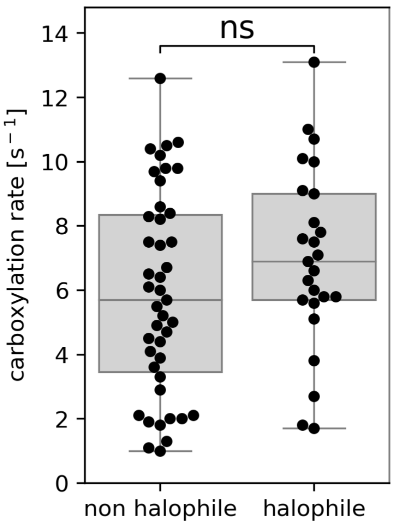


***Supplementary Fig. 7. Rubisco carboxylation rate is not significantly associated with host halotolerance.*** *Bacteria relations to environment salinity were collected from literature and plotted against rubisco carboxylation rates measured in vitro. Mann-Whitney U test was applied. ns, non significant.*

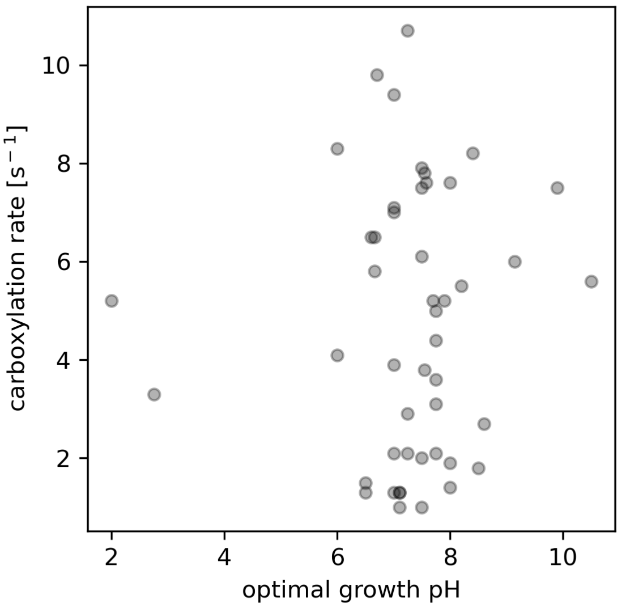


***Supplementary Fig. 8. Rubisco carboxylation rate is not significantly associated with the optimal growth pH in the host environment.*** *Bacteria optimal pH were collected from literature and plotted against their rubisco carboxylation rate measured in vitro.*

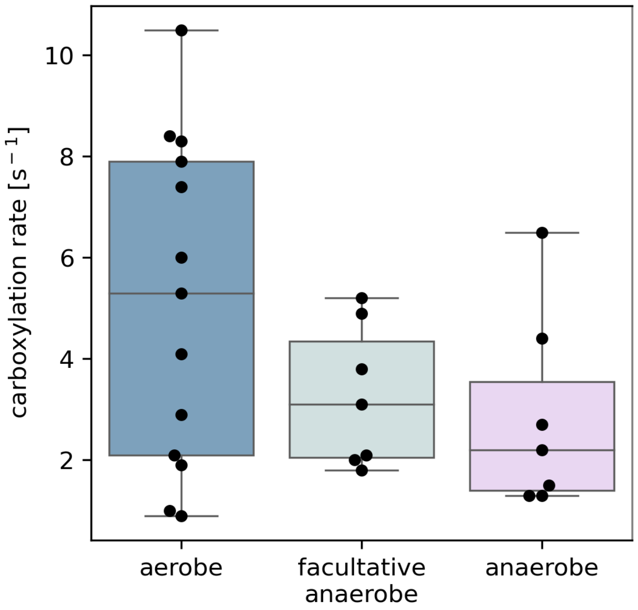


***Supplementary Fig. 9. Rubisco carboxylation rate across bacteria with different oxygen tolerance.*** *Bacteria relations to oxygen were collected from literature and plotted against rubisco carboxylation rates measured in vitro. Kruskal-Wallis test indicated no significant variation between the groups.*

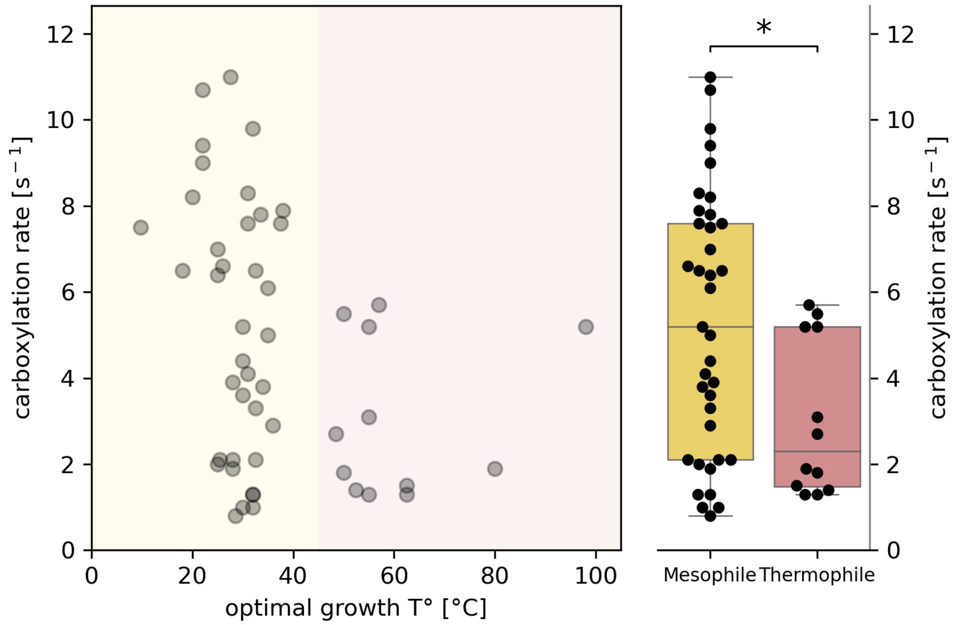


***Supplementary Fig. 10. Rubisco carboxylation rate as a function of the optimal growth temperature of its host bacteria.*** *Bacteria with optimal growth temperature below (yellow) and above (red) 45°C were considered as mesophiles and thermophiles respectively. Carboxylation rates of rubiscos from these 2 clusters were compared (right). Mann-Whitney U test was applied.* $^*p < 0.05$.

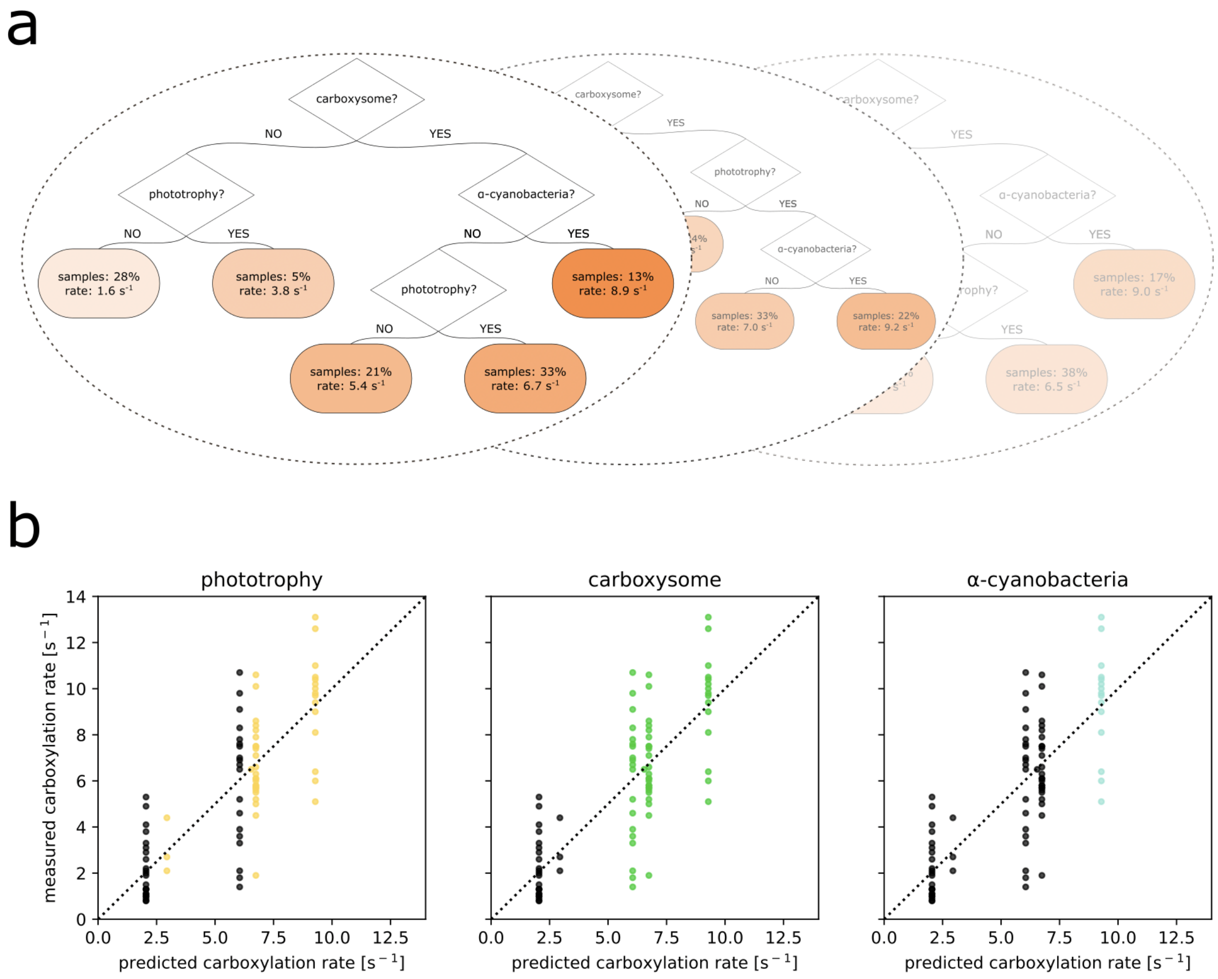


***Supplementary Fig. 11. Random forests modeling of form I rubisco carboxylation rate as a function of the main parameters from this study.*** *(A) Examples of decision trees out of the hundred trained in the random forests. (B) Measured against predicted carboxylation rates for the three most influential features according to the model. Each dot is color-coded based on the value of the respective feature (yellow/black: phototrophic/chemotrophic; green/black: carboxysome/non-carboxysome associated; cyan/black: α-cyanobacterial/non-α-cyanobacterial rubisco). RMSE = 2.1 $s^{-1}$; Average Explained Variance Score = 0.55.*

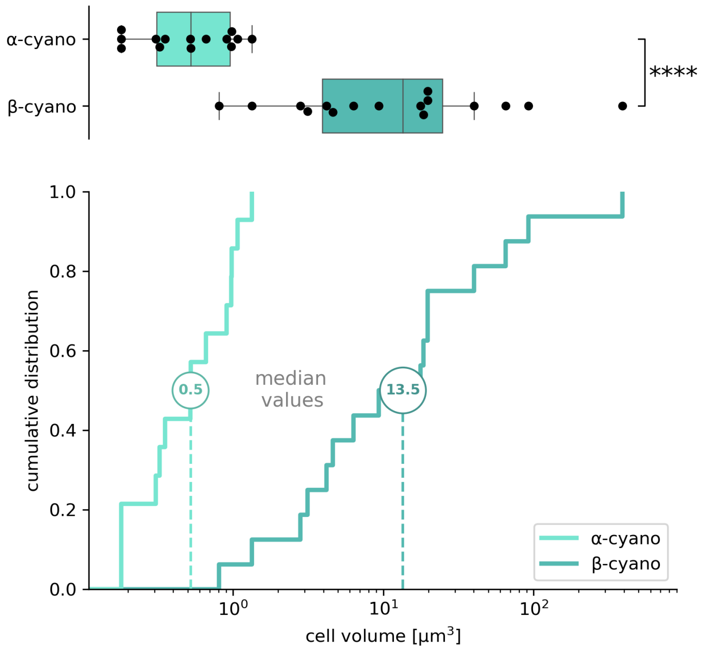


***Supplementary Fig. 12. α-cyanobacteria are smaller than β-cyanobacteria.*** *Box and cumulative distribution plots of the volume of α- and β- cyanobacterial cells. Size of cyanobacterial cells expressing rubiscos from Figure 2C were collected from the literature when possible and cell volume was calculated. Mann-Whitney U test was applied. ****p < 0.0001. Legend abbreviations are as follows: α-cyano, α-cyanobacteria; β-cyano, β-cyanobacteria.*

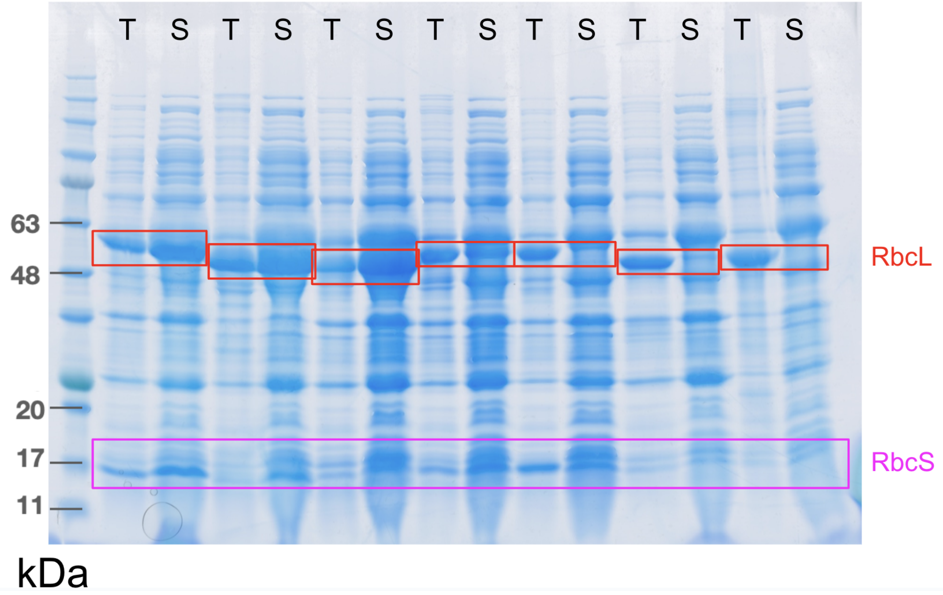


***Supplementary Fig. 13. SDS-page analysis of form I rubisco variants expression and solubility.*** *A characteristic gel of total (T) and soluble (S) extracts from E. coli cells expressing 7 different form I rubisco variants. Rubisco large (RbcL - red boxes) and small (RbcS - purple box) are ≈50 and ≈15 kDa monomers respectively; protein ladder is BLUeye prestained (GeneDirex hy-labs).*

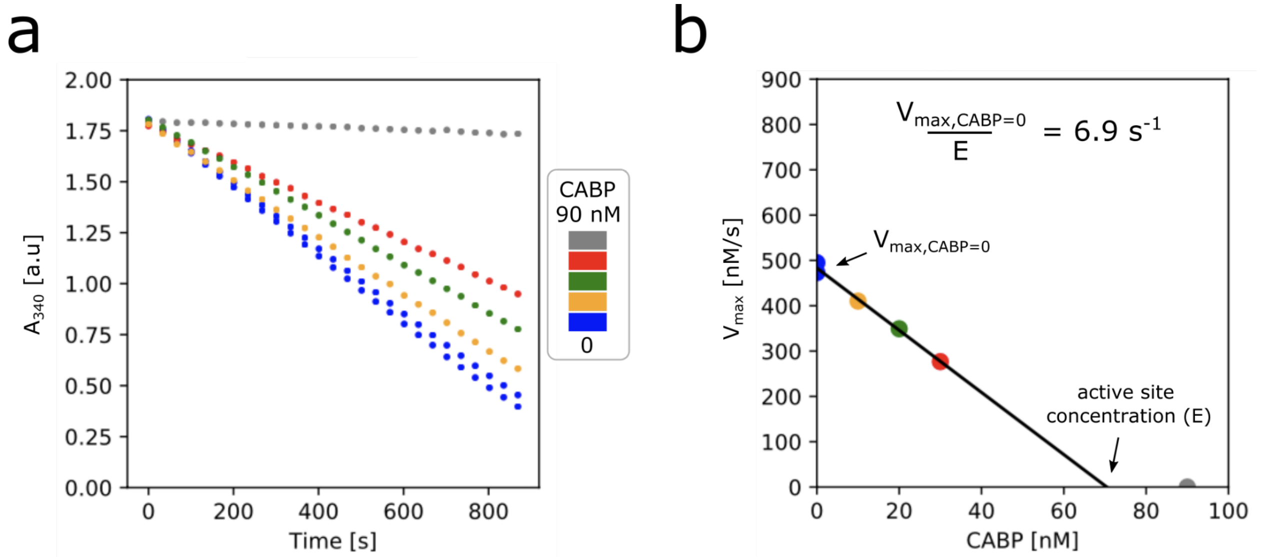


***Supplementary Fig. 14. Spectroscopic enzymatic assay for measuring the carboxylation rate of rubisco from a cellular extract (as presented in Davidi et al, 2020 (2)).*** *(A) Rubisco activity is coupled to NADH oxidation monitored at 340 nm. A gradient of CABP is used to gradually inhibit rubisco activity. Rubisco's maximal velocity ($V_{max}$) corresponds to half of the rate of NADH oxidation, which is the slope of the curves. (B) Rubisco's maximal velocity as a function of CABP concentration. The carboxylation rate of rubisco is given by dividing the maximal velocity without CABP (y-intercept) by the concentration of rubisco active sites (x-intercept). In this example, the carboxylation rate of R. rubrum (6.9 $s^{-1}$) is measured from a cellular extract without purification.*

***Supplementary Table 1. Recipe for the spectroscopic coupled assay.***

| Component | Assay concentration | Source |
|---|---|---|
| EPPS buffer pH 8.0 | 100 mM | Alfa Aesar (Cat # J61296) |
| MgCl2 | 20 mM | Sigma Aldrich (Cat # M2670-500G) |
| Dithiothreitol | 0.5 mM | Bio Basic Canada inc. (Cat # DB0058) |
| ATP | 2 mM | Sigma Aldrich (Cat # A3377-5G) |
| Phosphocreatine | 10 mM | Sigma Aldrich (Cat # 27920-5G) |
| NADH | 1.7 mM | Merck (Cat # 481913-1GM) |
| Carbonic anhydrase | 0.1 mg/mL | Sigma Aldrich (Cat # C3934-100MG) |
| Creatine phosphokinase | 20 U/mL | Sigma Aldrich (Cat # C3755-35KU) |
| Glyceraldehyde 3-phosphate dehydrogenase | 20 U/mL | Sigma Aldrich (Cat # G2267-10KU) |
| 3-Phosphoglyceric phosphokinase | 20 U/mL | Sigma Aldrich (Cat # P7634-5KU) |

### Supplementary Note 1: Rubisco expression and carboxylation assay from cell lysate

All 144 rubisco variants were cloned into a pET-29b vector and transformed into *E. coli* BL21 cells priorly transformed with a GroEL-GroES plasmid. Cells were grown to mid-log ($OD_{600}$ ≈ 0.6). After induction of GroEL-GroES and rubisco expression, cells were incubated overnight at 23°C. Upon cell harvest, the expression and solubility of each rubisco variant was assessed by SDS-page of cell extracts before (total lysate) and after (soluble extract) centrifugation (Supplementary Fig. 13).

Soluble extracts were then tested for carboxylation with a spectroscopic enzymatic assay. The assay was performed as described in Davidi *et al. (2)* Namely, all assay components (see Supplementary Table 1) were mixed and distributed into a 96-multiwell plate, except for the soluble cell extract and rubisco's substrate ribulose 1,5-bisphosphate (RuBP). After activation at 4% $CO_2$ and 0.4% $O_2$ for 15 minutes, soluble cell extracts were added to the wells and incubated for a further 15 minutes at the same gas conditions and at 30°C. The assay started upon the addition of RuBP to the wells and was immediately monitored for NADH oxidation (A340) in the gas-controlled plate reader. To measure the active site concentration, each variant was tested with 6 different concentrations of 2-C-carboxyarabinitol 1,5-bisphosphate (CABP) (0, 0, 10, 20, 30, and 90 nM) in parallel. CABP is a transition state analog and a stoichiometric rubisco inhibitor commonly used for active-site quantification (3). Rubisco's carboxylation rate is determined by measuring the slope of the linear regression fitted between the reaction rates and the CABP concentrations (Supplementary Fig. 14), allowing us to eliminate any background NADH oxidation reactions due to other native *E. coli* proteins.

Because it was not possible to estimate *a priori* the concentration of rubisco's active site in each extract, a first assay was performed with 10 µL of undiluted extract, often leading to saturation of this first assay. Following this, the concentration of each cell extract was adjusted through dilution to achieve a rubisco concentration that allowed measurable inhibition by CABP.

This pipeline enabled the determination of the carboxylation rate for ≈100 form I rubisco variants in a high-throughput manner.

### Supplementary Note 2: Effect of cell size on carbon uptake

Here we discuss the potential effect of cell size on carbon uptake by cyanobacteria. Existing literature suggests that cyanobacteria size is too small to have an effect on carbon uptake to the carboxysome (4, 5). However, these studies usually assume micron-sized cells, representing picocyanobacteria but not macrocyanobacteria (6). We reevaluate this logic with a back of the envelope calculation with variable cell size (while not including other effects such as the differences in the efficiency of inorganic carbon pumps etc.).

We consider a spherical cell with a radius R and a concentration of inorganic carbon ($C_i$) in the medium $[C_i]_{ext}$.

Using Fick's first law, we can determine the amount of $C_i$ taken by this cell per second. The equation is given by:

$\frac{dC_i}{dt} = -D_{C_i} \times S_r \times \frac{d[C_i]_r}{dr}$ (with $[C_i]_r$ the $C_i$ concentration at a distance, r, away from the center of the cell. And $S_r$ the surface of this r-radius sphere).

After differentiation, and with $[C_i]_r = [C_i]_{ext}(1 - \frac{R}{r})$ (considering $C_i$ null at r = R)

$$\frac{dC_i}{dt} = -D_{C_i} \times 4\,\pi\, r^2 \times (-\frac{R \times [C_i]_{ext}}{r^2})$$

$$\frac{dC_i}{dt} = D_{C_i}\, 4\,\pi\, R\, [C_i]_{ext}$$

The values used are as follows (7, 8):

- $D_{C_i} = D_{HCO_3-} = 10^{-9}\, m^2.s^{-1}$ (among all inorganic carbon forms, we consider $HCO_3^-$, the most abundant form, and the one going into carboxysomes)

- $[C_i]\ in\ the\ ocean \approx 10^{-3} M = 1\, mol.m^{-3}$

- $[C_i]\ in\ a\ freshwater\ lake \approx 10^{-4} M = 10^{-1}\, mol.m^{-3}$

- $R = 10^{-6}\, m$ (assuming a cell radius in the order of a micron)

So $\frac{dC_i}{dt} \approx 10^{-16}\, mol.s^{-1}\ (or\ 10^{10}\ C_i.s^{-1})$ in the ocean.

And $\frac{dC_i}{dt} \approx 10^{-15}\, mol.s^{-1}\ (or\ 10^{9}\ C_i.s^{-1})$ in a freshwater lake.

Theoretically, the minimum time to get all the carbon to divide is $t_{min} = \frac{n_{C\ in\ a\ cell}}{\frac{d[C_i]}{dt}}$

In order to make it more realistic, one should consider 1. the inherent limitations of diffusion from the medium to the carboxysome (as not all cell surfaces are equipped with carbon transporters and aspects such as $HCO_3^-$ diffusion and pool renewal may not be optimized in nature) and 2. the simultaneous loss of carbon atoms by cells through processes like respiration and fermentation, we propose a more realistic estimation by multiplying the minimum time by a factor of 10.

So, more probably, $t_{min} = 10 \times \frac{n_{C\ in\ a\ cell}}{\frac{d[C_i]}{dt}}$

For a cell with a radius of 1μm, the number of carbon atoms in the cell is $n_{C\ in\ a\ cell} \approx 10^{10}$ .

So $t_{min} = 10\ s$ for a cell with a radius of 1µm that thrives in the ocean and $100\ s$ in a freshwater lake.

Cyanobacteria have doubling times in the order of once a day. So, in both cases, $t_{min}$ << $T_d$ for a cell with a radius of 1 µm.

But some cyanobacteria (like *Limnoraphis robusta CS-951*) have a radius of 10 µm*.
For these cells, $n_{C\ in\ a\ cell} \approx 10^{13}$ and $\frac{dC_i}{dt} \approx 10^{11}\ C_i.s^{-1}$ in the ocean and $10^{10}\ C_i.s^{-1}$ in a freshwater lake.

So, for such cells, $t_{min} = 10^3\ s\ \approx\ 0.3\ h$ in the ocean and $3\ h$ in a freshwater lake.

We can therefore see that, in this last situation, the minimum time to get all the carbon to divide approaches the cell doubling time. This may be accentuated in cases where dissolved inorganic carbon concentrations are lower than typically measured for photic zones in oceanic or freshwater environments. In the case of α-cyanobacteria which were shown to dominate their beta counterparts not only in the oceans, but also in freshwater environments (9), their small size could therefore contribute to better supplying $CO_2$ to the rubisco. Finally, this smaller size (and increased cell surface-to-volume ratio) may potentially enhance the acquisition of other resources, including light. This, in turn, could further contribute to fueling the energy-consuming CCM and better concentrate $CO_2$ around rubisco.

*Actually, these cells are probably even worse carbon uptakers as they are stacked cylinders with a radius of 10 µm (and with a trichome's length going up to 5 mm)
